# An *in vitro* model maintaining taxon-specific functional activities of the gut microbiome

**DOI:** 10.1101/616656

**Authors:** Leyuan Li, Elias Abou-Samra, Zhibin Ning, Xu Zhang, Janice Mayne, Janet Wang, Kai Cheng, Krystal Walker, Alain Stintzi, Daniel Figeys

## Abstract

The gut microbiome is a new target for therapeutics. *In vitro* high-throughput culture models could provide time-and-cost saving solutions to discover microbiome responses to drugs. Unfortunately, there has been no report of *in vitro* models capable of maintaining functional and compositional profiles resembling the *in vivo* gut microbiome. Here, we developed and validated a high-throughput culturing model named Mipro to maintain individuals’ microbiomes. The Mipro model quintupled viable bacteria count while maintained the functional and compositional profiles of individuals’ gut microbiomes. Comparison of taxon-specific functions between pre -and-post culture microbiomes showed Pearson’s correlation coefficient *r* of 0.83 ± 0.03. Moreover, the Mipro model also exhibited a high degree of *in vitro – in vivo* correlation (Pearson’s *r* of 0.68 ± 0.09) in microbial responses to metformin in mice fed a high-fat diet. Mipro provides a highly simulated gut microbiome for high-throughput investigation of drug-microbiome interactions.

## Introduction

The gut microbiota is being increasingly recognized as a key factor in human health and disease ^1^ and as such, a target for drug therapy ^2^. Evidence is mounting that the gut microbiome can interact with drugs since the microbial metabolism can directly affect drug efficacy and toxicity ^3,4^, while drugs can alter the composition and function of the gut microbiota ^5–9^, which in turn impacts the host. Therefore, the gut microbial ecosystem, specific microbes, and microbial pathways are novel targets in drug discovery.

*In vitro* culture models could provide time-and-cost saving solutions to discover microbiome responses to drugs. However, current culture models do not maintain the functional and compositional profiles of the initial gut microbiome. To mimic an *in vivo* microbial ecosystem, it is key to develop culturing models and culturing media capable of conserving the composition and functional activities of an individual’s microbiome. Current *in vitro* culturing methods have been reported to sustain microbial diversity and achieve proper cultured bacterial coverage ^10–13^. However, we should not simply equal a microbiome’s diversity and coverage to its structure and functionality. Profound shifts in proportions of different taxa have been frequently described compared to the inoculum, in both batch culturing ^10,14^ and continuous flow culturing models, such as the Chemostat ^13^ and the SHIME ^12^. Importantly, shift of in microbiome composition can result in alternations of functional properties and ecological processes^15^, subsequently would result in different microbial responses to a drug stimulus. To the best of our knowledge, there has been no report on model maintaining microbiome properties and activities similar to the inoculum. In particular, preservation of functional activities hasn’t been described elsewhere. Moreover, comprehensive study of drug effects will require high-throughput screenings ^16^. Continuous flow models (e.g. Chemostat ^13^, SHIME ^12^ and M-SHIME ^17^) as well as microfluidic models (e.g. HuMiX ^18^, and gut-on-a-chip ^19^) cannot be readily adapted for high-throughput approaches, partially due to model size and the long period of time required for setting up and stabilization with these bioreactors.

For gaining a deep insight into drug responses of a microbiome, a technique that can precisely quantify microbial functional activities is required. Among the various meta-omics tools, metaproteomics directly quantifies the microbial functional responses at the end-product level, i.e. expressed proteins ^20^. The development and application of mass spectrometry (MS)-based metaproteomics technology in gut microbiome research has thrived in recent years. The identification coverage and sensitivity of MS-based metaproteomics has increased dramatically, enabling in-depth analysis of microbiome functional activities ^21,22^. Moreover, compared with sequencing-based techniques, metaproteomics has been validated to be more accurate for biomass estimates on a species level ^23^, and it also enables the study of taxon-specific functions through annotation of unique peptides ^24^. It is important that taxon-specific functional profiles in the microbiome is compared for revealing cultured stability and drug responses. This is because overall functions can maintain relatively stable due to the redundancy of functional genes among species in a gut microbiome ^25^. Therefore, metaproteomics is an adequate tool for insights into *in vitro* drug responses of gut microbiome.

Here, we report the development and validation of a high-throughput *in vitro* model for the <u>M</u>aintenance of gut microbiome <u>Pro</u>files (MiPro). Briefly, the MiPro model adopts an optimized culture medium and a 96-deep well plate-based format for microbiome culture (Figure 1A). The culture medium and culturing conditions were improved from our previous medium composition study ^26^. The model was first evaluated for its ability to maintain gut microbiome profiles *in vitro*, followed by testing and evaluation of the model’s *in vitro* correlation with *in vivo* drug response. This culture model enables, in combination with metaproteomic analysis, the assessment of drug effects on the microbiome at the compositional and functional levels. This model maintains high microbiome compositional stability and > 83% taxon-function similarity over 24 hr. In addition, we demonstrate that our MiPro model recapitulates the *in vivo* effects of metformin observed on individual mouse gut microbiomes.

**Figure 1.**
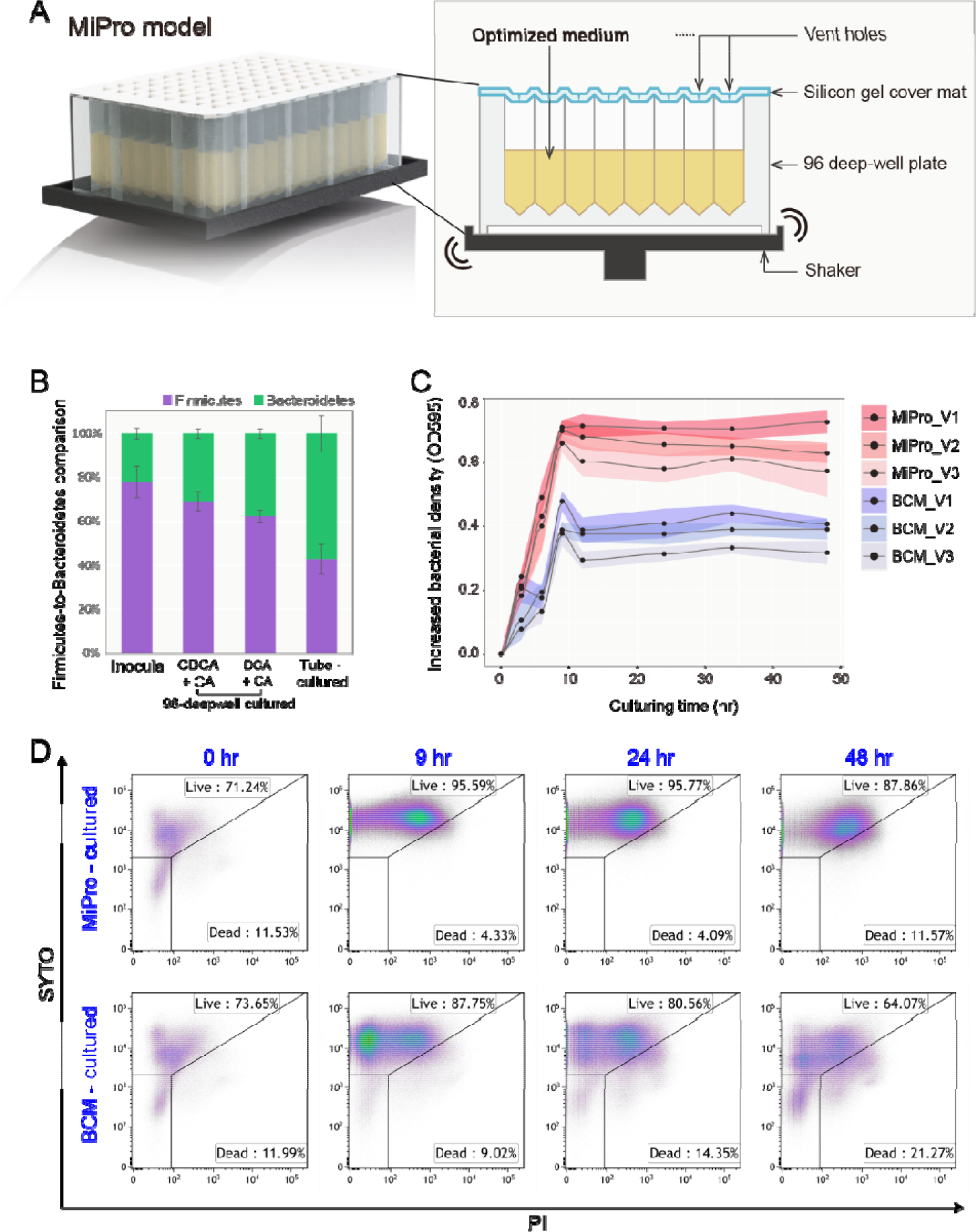
Establishment and general performance of the Mipro model. (A) Main components of the Mipro model: microbiome samples are cultured in an optimized culture medium in a 96-deep well plate. The plate is covered with a silicone-gel cover perforated at the top of each well. The plate is shaken at 500 rpm on a digital shaker. (B) Pre-experiment showing the proportion of Firmicutes and Bacteroidetes in the inocula (0 hr baseline sample), 96-deep well cultured microbiome with the presence of primary bile salts (CDCA and CA), 96-deep well cultured microbiome with the presence of commercialized bile salts mixture (DCA and CA), as well as tube-cultured microbiome with the presence of primary bile salts. All cultured samples were harvested and tested at 24 hr. Data shown are Mean ± SD (n = 3). (B) Increase in bacterial biomass over time in each individual microbiome (determination using absorbance at 595 nm). Colored ribbons indicate the range of standard deviations around the means (n = 4). (C) Temporal microbiome viability changes as shown with flow cytometry. The gating strategy is shown in Supplementary Figure S1.

## Results

### Establishment of the Mipro model

The first objective of this work was to develop a high-throughput *in vitro* model for the maintenance of gut microbiome profiles. We have recently evaluated the composition of the culture medium for optimal culturing of *ex vivo* microbiotas ^26^. Here, we improved upon the composition of our previously reported medium by assessing the effect of bile salts formula on the gut microbiome. Two formulae were compared: (1) a mixture of primary bile salts, i.e. 1:1 (w/w) sodium salt forms of cholic acid (CA) and chenodeoxycholic acid (CDCA), and (2) a commercialized 1:1 (w/w) mixture of sodium salt forms of CA and deoxycholic acid (DCA). The commercialized bile salts mixture has been adopted in a majority of gut microbiome culture media ^13,14,27–33^. The ratio of Firmicutes to Bacteroideteshas been extensively used to characterize microbiomes ^34^. Globally, the inoculum ratio of Firmicutes to Bacteroidetes better approximated to the ratio from the gut microbiota grown for 24 hr in the presence of CDCA and CA than that from the microbiome grown in presence of commercialized bile salts (Figure 1B).

Additionally, we assessed whether the gut microbiome culture was affected by the culturing conditions, namely (1) tube-based and (2) 96-deep well based culturing, while keeping all other conditions, including medium, inoculum, temperature, and container material (of polypropylene) constant. Culture tubes are the most frequently used containers in batch culture experiments ^14,26,33^. Notably, the 96-deep well was covered with a silicone-gel cover, which was perforated at the top of each well. This cover could prevent gas-exchange with the outer environment in the chamber, so as to preserve the partial pressure of gases and volatile metabolites in each well, which could subsequently preserve certain levels of dissolved gas molecules in the culture medium. In contrast to the use of 96-deep well plates, employing culture tubes resulted in a remarkable change in the proportions of Firmicutes and Bacteroidetes (Figure 1B). From the above results, we established our MiPro model (Figure 1A): a 96-deep well based culturing in combination with the optimized medium which contains 1:1 (w/w) of CA and CDCA (hereafter called MiPro medium).

### MiPro increased viable bacteria ratio and quintupled the count

We next evaluated the ability of the MiPro model to sustain a viable microbiome, which is vital for an effective *in vitro* microbiome response study. Temporal changes in parameters including microbial biomass, viability and diversity were compared with the 0 hr baseline sample. The commonly used basal culture medium (BCM) was included for comparison. At 24 hr the OD_595_ (bacterial intensity) was 0.8 ± 0.1 fold higher in the optimized medium as compared to that cultured in BCM (Figure 1C). Also, our MiPro medium achieved high ratio of viable bacterial cell throughout the culture period (95.77% at 24 hr compared with 71.24% at 0 hr, Figure 1D). We observed a 4.4-fold increase of viable bacterial count in MiPro medium after 24 hr of culturing, whereas a 3.0-fold maximum increase was detected in the BCM medium at 9 hr post-culturing.

### Mipro model maintains taxon-specific functional profiles of gut microbiome

In order to evaluate the effectiveness of MiPro to simulate the *in vivo* features of the gut microbiome inoculum, we used a metaproteomic approach ^35,36^ to characterize the taxonomic and functional stability of three individuals’ gut microbiomes over 9, 24, 34 and 48 hr of growth in either MiPro or BCM media. A minimum of three cultured, technical replicates were analyzed at each time point by LC-MS/MS. Ninety high-quality MS raw files were obtained with a total of 2,066,069 MS/MS spectra. A total of 58,848 peptides and 16,326 protein groups were identified with a false discovery rate (FDR) threshold of 1% (Supplementary Figure S2A). A high concordance (Pearson’s correlation coefficient *r* = 0.97 ± 0.02) was observed between the technical replicates of each group, indicating robust experimental reproducibility (Supplementary Figure S2B). Using an LCA approach on the MetaLab ^36^, a total of 21,839 peptides were assigned with a taxonomic lineage, resulting in 788 assigned species. The quantitative information (summed peptide intensities) was used to assess the species-level biomass contributions ^35,36^. 121 species that were quantified with ≥ 3 peptides were included in the comparison of species biomass contributions.

We applied a Bray-Curtis dissimilarity-based approach ^37^ for evaluating the variation of species biomass contributions between groups (Figure 2A). All MiPro-cultured microbiomes clustered closely with their corresponding inocula (0 hr baseline samples). In contrast, the BCM-cultured microbiomes were well separated from their inocula and exhibited greater dispersion over time than the MiPro-cultured microbiomes. Analysis of similarity (ANOSIM) based on the Bray-Curtis distance showed that at all time points, OM-cultured samples had an ANOSIM’s *R* of 0.122, *p* = 0.025; while the BCM cultured microbiome had an *R* of 0.2892, *p* = 0.001. Moreover, species-level biomass distribution bar-charts showed that the microbiome composition of three different individuals were sustained throughout the culturing period in the presence of MiPro medium (Figure 2B and Supplementary Figure S3). In particular, one-way ANOVA analysis on the latter dataset showed that the average relative abundance of *Faecalibacterium prausnitzii*, one of the most abundant gut microbial species quantified with metaproteomics in this dataset, significantly decreased in the BCM medium in comparison to both MiPro culture and baseline samples (Supplementary, Figure S4). In addition, both MiPro and BCM media achieved well-maintained alpha-diversity (Shannon-Wiener index) overtime across the three volunteers (Supplementary Figure S5).

**Figure 2.**
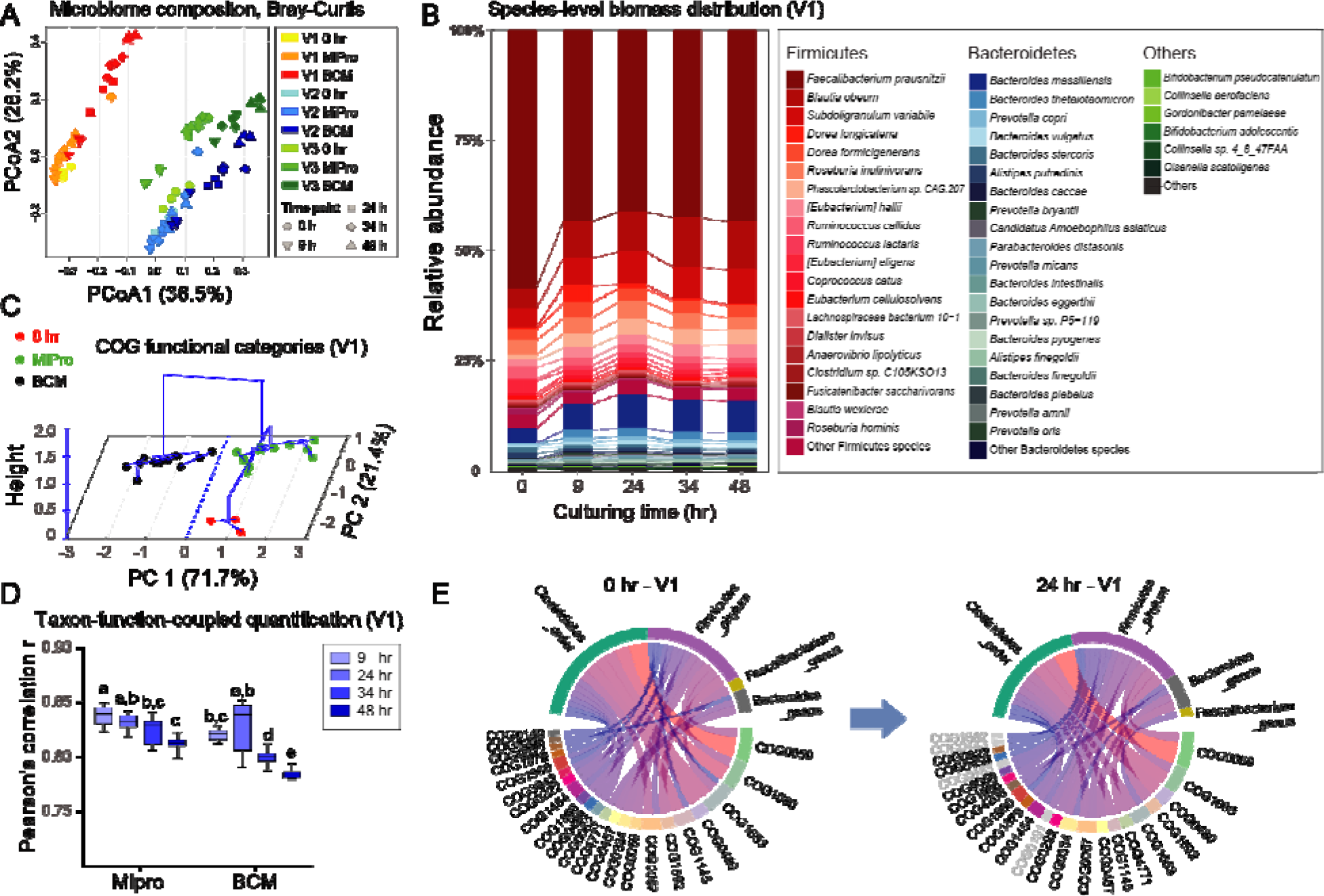
Metaproteomics revealed taxonomic & functional composition stability over time. (A) Principal coordinate analysis (PCoA) plot with Bray-Curtis dissimilarity on species level. (B) Compositional bar chart showing species-level biomass distribution over time in the cultured microbiome of V1 (see Supplementary Figure S3 for V2 and V3). (C) PCA scores plot with hierarchical clustering based on COG functional categories of individual V1 (see Supplementary Figure S6 for individuals V2 and V3). (D) Pearson’s correlation coefficient *r* of taxon-specific functional profiles between microbiome cultured over time and the baseline inoculum sample of V1 (see Supplementary Figure S7 for V2 and V3; different letters indicate significant differences at the *p* = 0.05 level, Tukey-b test; box spans interquartile range (25th to 75th percentile), and line within box denotes median. Whiskers represent min to max values.). (E) Visualization of associated taxa-functions enrichment analysis between 24 hr MiPro-cultured and 0 hr baseline samples in V1 (top 30 correlations were shown). Width of blocks and linkages represents number of matched proteins. Gray letters indicate un-paired pathways within the top 30 connections.

To assess the stability of functional activities, the identified proteins were annotated with clusters of orthologous group (COG) categories and the abundance of each COG category was calculated by summing the LFQ intensities of all the proteins belonging to the same COG category ^38^. Principal component analysis (PCA) was used to assess the relatedness of the samples based on the functional makeup of the metaproteomes. The first two components, PC1 and PC2, explained 71.7% and 21.4% of the total variance, respectively (Figure 2C). The largest functional variability was found in response to the culture condition using the BCM medium, indicating a better maintenance of the inoculum’s microbial functional profile by using the MiPro medium (Figure 2C and Supplementary Figure S6). In order to assess the maintenance of taxon-specific functional traits, we carried out a taxon-function-coupled analysis using the iMetaLab platform (http://shiny.imetalab.ca/) ^39^. In total we identified 1,066 unique COGs of proteins corresponding to 419 taxa, generating a three-dimensional dataset (sample-taxon-function) for between-sample comparisons. Pearson’s correlation coefficient *r* of the taxon-specific functional profiles was calculated between the inoculum and the cultured microbiome for each time point (Figure 2D). Statistical analysis based on Tukey’s b post hoc test of the changes in the *r*-value indicated that when compared with BCM, the MiPro culture achieved significantly higher taxon-specific functional correlations with baseline samples (*p* < 0.05; Figure 2D and Supplementary Figure S7). Pre-post culture correlations of taxon-function-coupled profiles reached an average of *r* = 0.83 ± 0.03 with our model (an average of all time points). To gain a more intuitive understanding of this correlation, we performed a taxon-function-coupled enrichment analysis using iMetaLab. Figure 2E shows the top 30 correlations (determined by the numbers of taxon-function matches) between the taxa and functions of the inoculum microbiome (0 hr baseline) and the 24 hr cultured microbiome for one individual. Similar taxa-function profiles were observed between the inoculum and 24 hr cultured microbiome indicating functional robustness in the MiPro assay.

### Evaluation of *in vitro-in vivo* correlation of microbial response to metformin treatment

We then evaluated the *in vitro-in vivo* correlation (IVIVC) of microbiome drug response in the MiPro model using high-fat diet (HFD) – fed C57/BL6 mice. Metformin is a widely prescribed drug for treating type 2 diabetes and it has been reported that in human 30% of the oral dose can be recovered in feces ^40^. Several studies have focused on the effect of metformin on gut microbiota composition and functions ^5,6,41^. For these reasons, we employed metformin to validate our MiPro model against *in vivo* studies by investigating the impact of metformin exposure on gut microbial communities in mice and in our model. Briefly, the MiPro model was inoculated with the stool microbiome from each mouse and cultured for 24 hr in presence or absence of metformin. Mice were then treated daily for 28 days with 300 mg/kg of metformin through gavage, and stool samples were collected at days 0, 14 and 28. All samples were then analyzed using metaproteomics. Thirty-four high-quality MS raw files were obtained with 926,004 MS/MS spectra. 51,294 peptide sequences corresponding to 12,733 protein groups were identified with an FDR threshold of 1%.

The IVIVCs of microbial response to metformin were evaluated for both taxon biomass contribution and functional activities. *In vitro* and *in vivo* variations in phylum-level biomass contributions were compared, as shown in Figure 3A. Increases in Bacteroidetes and decreases in Firmicutes were observed in both *in vitro* and *in vivo* treatments (Figure 3A). A high level of IVIVC at the phylum level, especially for day 14 samples, was indicated with Pearson’s correlation of *r* = 0.93 ± 0.02 (Figure 3B). Furthermore, at the genus level, we observed a consistent increase in the genera *Akkermansia*, *Bacteroides* and *Parabacteroides* (Figure 3C) under both *in vitro* and *in vivo* conditions, the *in vivo* change was agreement with the previously reported studies ^5,41,42^. At the species level, three separate groups of PLS-DA analyses were performed by comparing untreated microbiome versus microbiome following *in vivo* 14- and 28-day and *in vitro* 24 hr metformin treatment, respectively. Result showed that the biomass contribution of four bacterial species were consistently ranked with the highest VIP scores (VIP > 2 for all components) across all comparisons, and a 75% agreement in these changes was observed at the species level. Metformin-treatment increased the abundance of *A. municiphila*, which was in agreement with several previous studies ^42–45^.

**Figure 3.**
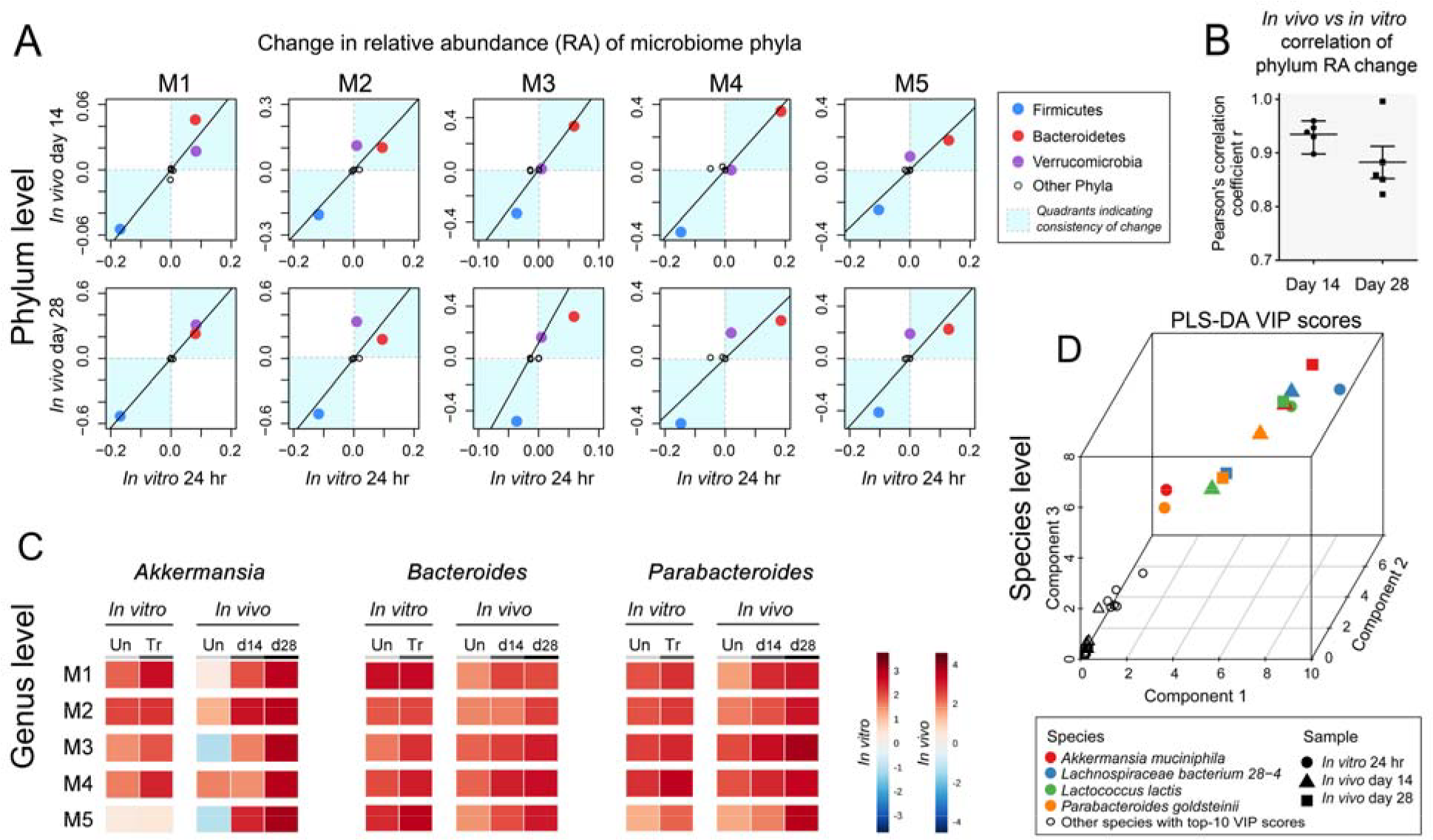
*In vitro – in vivo* correlation of taxonomic responses to metformin treatment. (A) Comparison of the change in relative abundance of the major gut bacterial phyla following *in vitro* and *in vivo* (days 14 and 28) metformin treatment of individual HFD-fed C57/BL6 wild-type litter-mate mice. Data point that falls into quadrants I and III (in blue) indicates *in vitro*-*in vivo* consistency in increased and decreased phyla in response to metformin, respectively. (B) Pearson’s correlation coefficient *r* between *in vivo* and *in vitro* changes of phyla in response to metformin treatment. (C) Heat map showing change in relative abundance of three bacterial genera, known to increase in metformin treated HFD-fed mice. The left panels represent *in vitro* cultured individual mouse microbiome in the absence or presence of metformin, whereas the right panels correspond to *in vivo* microbiome drug-treatment at days 14 and 28. (n = 5). (D) Comparison of variable importance in projection (VIP) on the first three components from three separate PLS-DA analyses. PLS-DA analyses were performed by comparing untreated microbiome vs microbiome following *in vivo* 14-day, 28-day and *in vitro* 24 hr metformin treatment, respectively. Four bacterial species were consistently ranked with the highest VIP scores in the three analyses.

For functional analysis, protein groups were first filtered with the criteria that the protein groups should be present in ≥ 50% in each of the listed subgroups (including *in vitro* untreated vs. *in vitro* 24 hr treated, *in vivo* untreated vs. *in vivo* day 14 or day 28 treated samples). Partial least squares discriminant analysis (PLS-DA) was performed on shared proteins, and proteins with a VIP score > 1 were regarded as differential protein groups, which were mainly involved in 12 KEGG pathways (Figure 4A). A total of 11,222 (out of 17,646) proteins that correspond to these KEGG pathways were extracted from the original protein group file. All LFQ intensities of protein groups that were assigned to each of the 12 pathways were summed in each sample for a KEGG pathway level evaluation. Figure 4A shows that the relative abundances of selected pathways were uniformly altered in both *in vitro* and *in vivo* metformin treated samples in comparison to untreated microbiomes. Subsequently, IVIVC were visualized by comparing the changed value of these KEGG pathways (Figure 4B). In most cases, changed proteins appeared in Quadrants I and III suggesting agreement of *in vitro-in vivo* responses. Among these, glycolysis/glucogenesis, ABC transporters and two-component system showed high levels of variation. Here we took a deeper look into the shift of functional balance by normalizing the intensities of each enzyme against the summed protein intensity of the glycolysis/gluconeogenesis pathway (Figure 4C). In general, it was observed that the proportions of enzymes under *in vitro* and *in vivo* were similar, suggesting well-maintained functional ecology for the *in vitro* model. Nine out of the 17 enzymes were significantly affected by metformin treatment *in vivo*; while for the *in vitro* model, five significant changes were found. All others enzymes, although not statistical significance, showed the same trends within the pathway.

**Figure 4.**
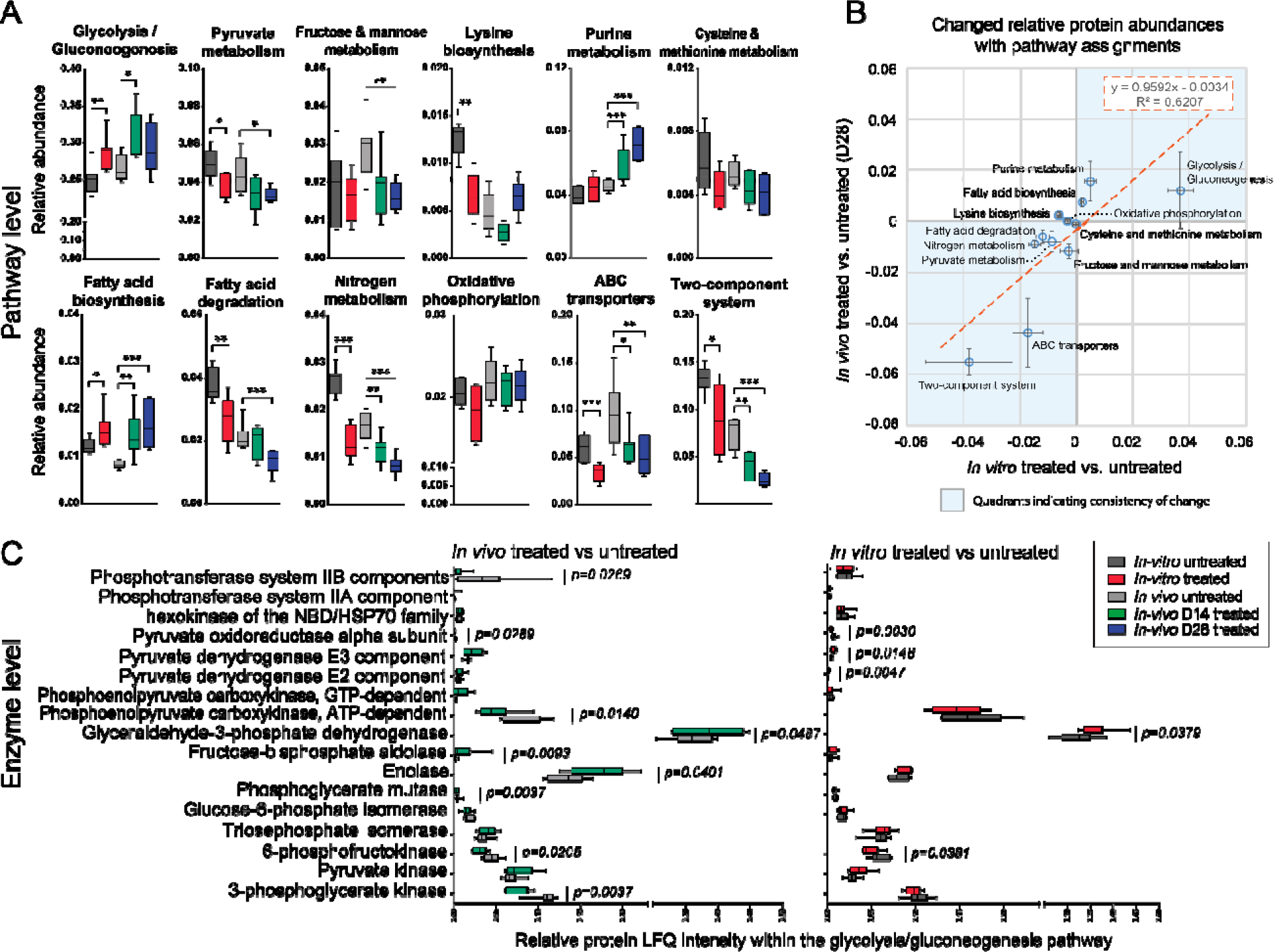
*In vitro – in vivo* correlation of microbiome pathway and enzyme responses to metformin. (A) Pathways analysis using the relative abundance of protein groups that are assigned to the selected KEGG pathways (KEGG pathways that corresponding to PLS-DA VIP > 1 protein groups). (B) *In vitro – in vivo* correlation of changed relative protein abundance with pathway assignments in microbiome samples following metformin treatment from HFD-fed mice (data shown as mean ± SEM, n = 7). (C) Detailed view of multiple enzymes abundance in the glycolysis/gluconeogenesis pathway. Data shown here represent the relative abundance of each enzyme normalized against the total intensity of all enzymes detected in the pathway. Levels of significance are according to two sided, non-parametric *t-test*: * *p* < 0.05, ** *p* < 0.01, *** *p* < 0.001. Box spans interquartile range (25th to 75th percentile), and line within box denotes median. Whiskers represent min to max values.

## Discussions

In order to build an *in vitro* model to assess microbiome function or to address microbiome’s response to xenobiotics, it is critical to preserve both the compositional and functional profiles of an individual’s microbiome. The gut microbiome constitute a complex microbial ecosystem and each microbe may play different functional roles ^46^. Environmental changes can trigger structural and functional alternations in the microbiome. Although some studies have shown the culturability of the gut bacterial community *in vitro* ^10,13,14,47^, these studies have not demonstrated the maintenance of the microbiota’s functional activities. While taxonomic profiling is important to characterize an individual’s microbiome, biological processes and functions cannot be assessed by compositional analyses. Yet, such information is essential for gaining deeper understanding of the microbiota’s behavior and functional activities and ultimately for designing microbiome-targeted manipulations. The need for functional understanding is even more crucial given that the microbiome’s functions can be altered in absence of compositional changes. For example, sialylated milk oligosaccharides elicit a microbiota-dependent growth of mice that are used as models of infant undernutrition by inducing the production of microbial metabolites, yet without any significant change in the relative abundance of most microbes ^48^. Elucidating the microbiome’s functions requires a comprehensive quantification of the transcripts, proteins and/or metabolites produced by the microbial community. Here we used a metaproteomic approach we recently developed ^38^ to simultaneously characterize the microbiome’s taxonomic biomass contributions and its functional activity while developing and validating a high-throughput *in vitro* model for the <u>M</u>aintenance of gut microbiome <u>Pro</u>files (MiPro).

In order to build an *in vitro* model to assess microbiome function or to address microbiome’s response to xenobiotics, it is critical to preserve both the compositional and functional profiles of an individual’s microbiome. Although some studies have shown the culturability of the gut bacterial community *in vitro* ^10,13,14,47^, these studies have not demonstrated the maintenance of the microbiota’s functional activities. The majority of gut microbiome culture media (including BCM) ^13,14,27–33^ contain a mixture of commercialized bile salts constituted of the sodium salt forms of CA and DCA at a 1:1 ratio (w/w). DCA (secondary bile acid) is known to act as an antimicrobial agent due to its high-hydrophobic and detergent properties on bacterial membranes ^49^. Studies have noted a decrease in Firmicutes abundance in response to DCA ^50^, whereas an increased level of primary bile acids drove Firmicutes’ enrichment ^51^. Hence, we replaced DCA with CDCA, a major primary bile acid produced in human ^52^. This replacement was effective in maintaining the *in vitro* microbiome composition (Figure 2A). In terms of culturing condition, a silicon-gel cover was perforated at the top of each well in order to create tiny vent holes permitting the escape of the gas produced by the gut microbiota while minimizing inward gas diffusion from the anaerobic chamber. We found that culture tubes with loosen caps resulted in marked microbiome composition changes, the 96-well format maintained the microbiome composition (Figure 2B). The gut microbiome produces gases such as H_2_, CH_4_, H_2_S and NO_*x*_, etc. ^53^ and volatile organic compounds, such as SCFAs ^54^. As opposed to culturing in a tube, the silicon-gel cover increased the partial pressure of gases and volatile metabolites in each well, which subsequently preserved certain levels of dissolved gas molecules in the culture medium. Some dissolved metabolites are important factors for bacteria cross-feeding ^54^ and the control of pathogens ^55^, and thus, are presumably essential for the maintenance of an *in vitro* microbiome. Besides the application in our high-throughput model, these optimizations may also be adoptable in previously-reported fluidic-based models ^17–19,56^, in which the optimized medium would help to maintain microbial functional activities, and a similar action that preserves a proper gas metabolite pressure should be considered.

As an outcome of these optimizations, our MiPro model quintupled viable bacteria count, and had similar alpha-diversity and microbial composition at 24 hr post-inoculation to the inoculum at time 0. These findings are in agreement with the premise that high biodiversity may enhance the temporal stability of microbial communities ^57^. In terms of functions, a general view of functional stability using summed protein abundances in each functional category is not sufficient due to the complexity of functional constitution in a gut microbiome ecosystem. Study has shown that at the microbiome level, functional pathways are stable within a health human population ^58^. This is stability owes to the functional redundancy among species in a gut microbiome ^25^. Functional compensation can happen among different taxa, which preserves long-term average ecosystem performance, as one species can increase a function for loss or decline in another ^59^. This mechanism shifts the functional balance between two taxa in a gut ecosystem. Therefore, for maintenance of an *in vitro* gut microbiome culturing system, it is very important to preserve the stability of taxon-specific functional profiles in the microbiome. And results showed that this has been well achieved in our model. Together, our MiPro model comprises two phases, a growth phase with significant viable biomass accumulation and a lag phase where > 83% of the taxon-specific functional activities of the inoculum are maintained. Hence, this model can be effectively used to amplify a microbial community and to investigate the effect of perturbations on the microbial community composition and functional activities.

The IVIVC study estimated the power of the MiPro to recapticulate *in vivo* drug responses on different levels of taxonomic biomass contributions and functional activities. Changes of major taxonomic responders on phylum, genus and species levels as reported in several studies ^5,41–45^, were captured in both our *in vitro* and *in vivo* treatments. Additionally, our functional profiling analysis showed agreements of *in vitro – in vivo* responses, and as well as with previous studies ^5^. significant responses were found in pathways of glycolysis/gluconegenesis, pyruvate metabolism, fatty acid biosynthesis, fatty acid degradation, nitrogen metabolism, ABC transporters, and the two-component system. A recent study ^5^ indicated that metformin alters multiple microbial pathways including ABC transporters, two-component system, fructose and mannose metabolism and pyruvate metabolism. All of these responses were found in our model. Two microbial ABC transporters and the two-component systems were significantly decreased, while the glycolysis/ gluconeogenesis pathway was similarly increased following drug treatments in both the MiPro and *in vivo* models (Figure 5C). Shin *et al.* has reported the role of metformin in improving glucose homeostasis in HFD-fed mice ^42^. Changes in glycolysis/gluconeogenesis pathway inside the gut microbial environment could play a key role in glucose absorption across the intestinal mucosal layer ^60^. Notably, metaproteomic responses within the glycolysis/gluconeogenesis pathway revealed high IVIVC of the Mipro on the enzyme level (Figure 4C). This highlighted the depth of our model to recapitulate *in vivo* microbiome functional activities in response to drug treatment.

## Conclusions

In this work, we evaluated the performance of the MiPro model to maintain microbial taxon-function stability and the utility of this model for drug-microbiome interactions studies. We optimized the medium and culture model for high-throughput drug-microbiome co-culturing. The optimized model showed improved performance in sustaining viability, diversity, compositional and functional profiles of the inoculum microbiome. A high degree of *in vitro – in vivo* correlation of compositional and functional responses were observed with metformin treatment. Our work provides an effective experimental platform for drug-microbiome interaction studies.

## Methods

### Medium preparation and gut microbiome culturing

The medium composition for MiPro was based on our previously suggested medium composition ^26^, which comprises: 2.0 g L^−1^ peptone water, 2.0 g L^−1^ yeast extract, 0.5 g L^−1^ L-cysteine hydrochloride, 2 mL L^−1^ Tween 80, 5 mg L^−1^ hemin, 10 μL L^−1^ vitamin K1, 1.0 g L^−1^ NaCl, 0.4 g L^−1^ K_2_HPO_4_, 0.4 g L^−1^ KH_2_PO_4_, 0.1 g L^−1^ MgSO_4_⋅7H_2_O, 0.1 g L^−1^ CaCl_2_⋅2H_2_O, 4.0 g L^−1^ NaHCO_3_, 4.0 g L^−1^ porcine gastric mucin (cat# M1778, Sigma-Aldrich), and 0.5 g L^−1^ bile salts. Based on this composition, two media containing either commercialized mixed bile salts (0.5 g L^−1^, 1:1 sodium cholate: sodium deoxycholate, cat# 48305, Sigma-Aldrich) or a primary bile salts composition (0.25 g L^−1^ sodium cholate and 0.25 g L^−1^ sodium chenodeoxycholate) were compared for further optimization. To test and improve the physical culturing condition, samples were cultured either in 1 ml of media in 96-deep well plates or in 2 ml of media in culturing tubes (cat#T406-2A, Simport, Canada). The 96-deep well plate was covered with a silicone gel mat with a vent hole on each well made by a sterile syringe needle. During culturing, plates and tubes were shaken at 500 rpm with digital shakers (MS3, IKA, Germany). The optimal medium and culturing condition for the MiPro model was then determined based on its ability to maintain gut microbiome composition assessed as described below.

Stool samples were collected in compliance with our Research Ethics Board protocol (# 20160585-01H) approved by the Ottawa Health Science Network Research Ethics Board at the Ottawa Hospital. Briefly, ~ 3 g of fresh stool sample was collected from each individual with a 2.5 ml sterile sampling spoon (Bel-Art, United States). The spoon was dropped into a 50 ml Falcon tube containing 15 ml of sterile PBS pre-reduced with 0.1% (w/v) L-cysteine hydrochloride. The samples were immediately transferred into an anaerobic workstation (5% H_2_, 5% CO_2_, and 90% N_2_ at 37°C). Before homogenization with a vortex mixer, the tube was uncapped for a few seconds to enable gas exchange and in particular oxygen removal. Sample homogenates were filtered using sterile gauzes and were immediately inoculated into each medium for static culturing at a final inoculum concentration of 2% (w/v). Culturing of an individual’s microbiome was carried out in the MiPro model containing 1 ml of the optimized culture medium. The conventional basal culture medium (BCM) ^14^ was also included for evaluating performance of our improved medium. For each individual, 32 replicates were cultured for each medium, allowing for 4 replicates at 8 different time points. Microbiomes were characterized at 0 (immediately after inoculation), 3, 6, 9, 12, 24, 34 and 48 hr by measurements of the optical density (OD) at 595 nm (as a proxy of microbial growth and biomass) and by metaproteomic analyses.

### Microbiome growth and viability tests

At 0, 3, 6, 9, 12, 24, 34 and 48 hr, two 100 μl aliquots were removed from each sample for OD_595nm_ measurements. One of the aliquots was centrifuged at 16,000 × g and the supernatant was used as the medium blank for the OD measurement. Viability of the microbiomes were tested at 0, 9, 24 and 48 hr using the LIVE/DEAD BacLight Kit (Thermo Fisher Scientific Cat# L7012) in combination with flow cytometry ^61^. Briefly, according to the manufacturer’s instruction, microbial cells were washed and diluted in 0.85% NaCl saline buffer. Then, the bacteria were stained with PI and SYTO for 15 minutes prior to acquisition on a BD FACSCelesta™ multicolor cell analyzer. For maximum bacterial cell viability, all sample processing procedures, including mixing, dilution and staining steps were performed inside the anaerobic station. Bacterial cells were gated according to size and granularity on the FSC/SSC scatter plot. Live bacterial count was done in reference to the SYTO^+^ PI^−^ gate. Data were analyzed using Kaluza Analysis Software version 1.5.

### MiPro model to *in vivo* comparison and validation

*In vitro* and *in vivo* effects of metformin on the murine gut microbiome associated with a high-fat diet (HFD) were tested. Briefly, 7-week old male litter-mates from inbred C57/BL6 mice were single-housed and fed a 42% fat calories diet (ENVIGO, TD.09682) for 6 weeks to allow stabilization of their microbiomes on diet. Fresh stool pellets from each mouse were collected for *in vitro* culturing on day-0. The microbiomes were cultured in the absence or presence of metformin (cat# PHR1084, Sigma-Aldrich) using the MiPro model. The *in vitro* concentration of metformin was set to 6 mg/ml, emulating the 30% fecal recovery ratio of metformin previously reported from *in vivo* experiments ^40^. Cultured microbiome samples were harvested at 24 hr for metaproteomic analysis. For the *in vivo* experiment, mice were treated daily with 300 mg/kg of metformin through oral gavage. Stool pellets were collected for metaproteomic analysis at days 0, 14 and 28. The animal experiment was performed at the University of Ottawa and conducted in strict accordance with the guidelines of the Care and Use of Experimental Animals of Canadian Council on Animal Care (CCAC). The animal use protocol BMI 2848 was approved by the Animal Care Committee at the University of Ottawa.

### Trypsin digestion, desalting and LC-MS/MS analysis

Protein extraction, trypsin digestion, and desalting steps were carried out as previously described ^38^, with a minor modification of the cell washing step. Briefly, for general analysis, the samples were washed two times with PBS at 14,000 × *g*, 4°C for 20 min. Then large debris were removed with 300 × g centrifugation at 4°C for 5 min, followed by pelleting cells at 14,000 × *g*, 4°C for 20 min. The microbial cell pellets were lysed by ultrasonication in 200 µL lysis buffer (4 % (w/v) sodium dodecyl sulfate and 8 M urea in 50 mM Tris-HCl buffer, pH 8.0; plus Roche PhosSTOP™ and Roche cOmplete™ Mini tablets), and proteins precipitated overnight in acidified acetone/ethanol at −20°C. Proteins were washed three times with ice-cold acetone, then dissolved in 6 M urea in 50 mM ammonium bicarbonate (pH 8). Protein concentrations were determined by DC (detergent compatible) protein assay before an overnight trypsin digestion at 37°C following protein reduction and alkylation, as previously described [25]. Desalted tryptic peptides corresponding to 1 μg of protein were loaded for LC-MS/MS analysis with an Agilent 1100 Capillary LC system (Agilent Technologies, San Jose, CA) and run on a Q Exactive mass spectrometer (ThermoFisher Scientific Inc.). Peptides were separated with a tip column (75 μm i.d. × 50 cm) packed with reverse phase beads (1.9 μm/120 Å ReproSil-Pur C18 resin, Dr. Maisch GmbH, Ammerbuch, Germany). Peptide separation was performed using either a 90 min gradient for human samples or a 120 min gradient for mouse samples. The gradients included 5 to 30% (v/v) acetonitrile at a flow rate of 200 nL/min, for which 0.1% (v/v) formic acid in water was used as solvent A, and 0.1% FA in 80% acetonitrile was used as solvent B. The instrument parameters included a full MS scan from 300 to 1800 m/z, followed by data-dependent MS/MS scan of the 12 most intense ions, a dynamic exclusion repeat count of two, and repeat exclusion duration of 30 s. All samples were run on LC-MS/MS in a random order. In addition, for the pre-experiment that evaluated bile salts composition and culture conditions, samples were analyzed on an Orbitrap XL following a 6 hr gradient using previously described parameters ^26^. All raw data from LC-MS/MS have been deposited to the ProteomeXchange Consortium (http://www.proteomexchange.org) via the PRIDE partner repository.

### Metaproteomics data processing

Protein/peptide identification and quantification were carried out using the MetaLab software (version 1.0) ^36^, which automates the MetaPro-IQ approach ^38^. An iterative database search strategy using gut microbial gene catalogs (for cultured human microbiomes, human gut microbial gene catalog with 9,878,647 sequences from http://meta.genomics.cn/ and for mice microbiomes, mouse gut microbial gene catalog database comprising 2,572,074 genes, obtained from http://gigadb.org) and a spectral clustering strategy were used for database construction, the peptide and protein lists were generated by applying strict filtering based on a FDR of 0.01, and quantitative information of proteins were obtained with the maxLFQ algorithm; quantitative taxonomic analyses were achieved by assigning identified peptides with taxonomic lineage of lowest common ancestor (LCA) through the pep2tax database and summing up the intensities of the peptides. Functional annotations (COG, KEGG) were obtained with MetaLab version 1.1.0.

For human gut microbiomes, protein groups were filtered with the criteria that the protein should be present in ≥ 25% of the samples (Q25). Bray-Curtis dissimilarity and analysis of similarities (ANOSIM) were performed using the R package “vegan”. Principal coordinates analysis (PCoA), principle component analysis (PCA), and hierarchical clustering were visualized with R (version 3.4.3). Taxonomic composition visualization and taxon-function coupled analysis were performed and visualized using iMetaLab (http://imetalab.ca/). The database of clusters of orthologous groups (COG) of proteins was used for functional annotation. For each sample, taxon-specific functional proteins with protein intensity was then generated from iMetaLab. With these set of tables, the Pearson’s correlation coefficient *r* of the taxon-function coupled profile between any two samples was calculated using R to generate a correlation matrix, and the correlation between 0 hr and the cultured microbiome at each subsequent time point was obtained. For visualizing the taxon-function coupled enrichment, *p* < 0.05 was set as the threshold for both taxonomic and functional enrichment, and the top 30 connections were selected from enriched taxon-function matches.

For the *in vitro-in vivo* mice microbiome experiments, protein groups were filtered with the criteria that the protein groups should be present in ≥ 50% (Q50) in each of the listed subgroups (including *in vitro* untreated vs. *in vitro* 24 hr treated, *in vivo* untreated vs. *in vivo* day 14 or day 28 treated samples), partial least squares – discriminant analyses (PLS-DA) test was performed on shared proteins among listed subgroups using the online tool MetaboAnalyst (www.metaboanalyst.ca/). Protein groups with a VIP score > 1 were annotated to the corresponding KEGG categories. Then all protein groups annotated with these KEGG categories were extracted from the original protein group file. *In vitro – in vivo* correlation of the microbiome drug response was evaluated using taxonomic and pathway change of metformin-treated microbiome. Pearson’s correlation coefficient *r* was calculated using R. Taxonomic composition analysis was done using MetaboAnalyst.

## List of abbreviations

ANOVA: analysis of variance
BCM: basal culture medium
CA: cholic acid
CDCA: chenodeoxycholic acid
COG: Clusters of Orthologous Groups
DCA: deoxycholic acid
FDR: false discovery rate
FSC: forward scatter
KEGG: Kyoto Encyclopedia of Genes and Genomes
KO: KEGG Orthology
LC-MS/MS: liquid chromatography-tandem mass spectrometry
LCA: lowest common ancestor
LFQ: lable-free quantification
MiPro: Maintenance of gut microbiome Profiles
OD: optical density
PCA: principle component analysis
PCoA: principal coordinates analysis
PLS-DA: partial least squares – discriminant analysis
SSC: sideward scatter

## Declarations

### Ethics approval and consent to participate

The human stool sampling protocol was approved by the Ottawa Health Science Network Research Ethics Board at the Ottawa Hospital (# 20160585-01H). All participants signed informed consent to participate in the study. The animal use protocol was approved by the Animal Care Committee at the University of Ottawa (# BMI 2848).

### Availability of data and material

All raw data from LC-MS/MS have been deposited to the ProteomeXchange Consortium (http://www.proteomexchange.org) via the PRIDE partner repository.

### Authors’ contributions

LL, EAS and DF designed the study. DF supervised the study. LL, EAS, JM, ZN, JW and KW performed the experiments. LL, EAS, ZN, XZ and KC performed data analysis. LL, EAS and DF wrote the manuscript. AS, DF, XZ, JM and ZN contributed to the editing and revision of the manuscript. All authors read and approved the final manuscript.

## Acknowledgements

DF acknowledges a Canada Research Chair in Proteomics and Systems Biology.

## Electronic supplementary material

Figure S1. Gating of live, dead and unstained bacteria according to stained gut microbiome cells, stained and heat-treated microbiome cells, and unstained microbiome. Figure S2. Metaproteomic data quality.

Figure S3. Compositional bar chart showing species-level biomass distribution over time in the cultured microbiome of volunteers V2 and V3. Figure S4. Comparison of *Faecalibacterium Praunsnitzii* biomass change in the MiPro- and BCM-cultured microbiomes. Figure S5. Shannon-Weiner index suggesting well-maintained alpha-diversity of the microbiomes cultured from volunteers V1-3 over 48 hrs. Figure S6. PCA scores plot with hierarchical clustering based on COG functional categories of microbiome proteins from volunteers V2 and V3. Figure S7. Taxon-function-coupled profile in comparison with 0 hr baseline samples.

## Reference

1. Kåhrström, C.T., Pariente, N. & Weiss, U. Intestinal microbiota in health and disease. Nature 535, 47 (2016).

2. Cani, P.D. & Delzenne, N.M. The gut microbiome as therapeutic target. Pharmacol. Ther. 130, 202–212 (2011).

3. Wilson, I.D. & Nicholson, J.K. Gut microbiome interactions with drug metabolism, efficacy, and toxicity. Transl Res 179, 204–222 (2017).

4. Hooper, L.V. & Gordon, J.I. Commensal host-bacterial relationships in the gut. Science 292, 1115 (2001).

5. Wu, H., et al. Metformin alters the gut microbiome of individuals with treatment-naive type 2 diabetes, contributing to the therapeutic effects of the drug. Nat Med 23, 850 (2017).

6. Zhang, X., et al. Modulation of gut microbiota by berberine and metformin during the treatment of high-fat diet-induced obesity in rats. Sci Rep 5, 14405 (2015).

7. Maccaferri, S., et al. Rifaximin modulates the colonic microbiota of patients with Crohn’s disease: an *in vitro* approach using a continuous culture colonic model system. J Antimicrob Chemother 65, 2556 (2010).

8. Xu, D., et al. Rifaximin alters intestinal bacteria and prevents stress-induced gut inflammation and visceral hyperalgesia in rats. Gastroenterology 146, 484 (2014).

9. Morgan, A.P., et al. The antipsychotic olanzapine interacts with the gut microbiome to cause weight gain in mouse. PLOS ONE 9, e115225 (2014).

10. Kim, B.-S., Kim, J.N. & Cerniglia, C.E. *In vitro* culture conditions for maintaining a complex population of human gastrointestinal tract microbiota. J Biomed Biotechnol 2011, 838040 (2011).

11. Lau, J.T., et al. Capturing the diversity of the human gut microbiota through culture-enriched molecular profiling. Genome Med 8, 72 (2016).

12. Kim, B.-S., Kim, J.N. & Cerniglia, C.E. *In vitro* culture conditions for maintaining a complex population of human gastrointestinal tract microbiota. Journal of Biomedicine and Biotechnology 2011, 10 (2011).

13. McDonald, J.A.K., et al. Evaluation of microbial community reproducibility, stability and composition in a human distal gut chemostat model. J Microbiol Methods 95, 167 (2013).

14. Long, W., et al. Differential responses of gut microbiota to the same prebiotic formula in oligotrophic and eutrophic batch fermentation systems. Sci Rep 5, 13469 (2015).

15. Costello, E.K., Stagaman, K., Dethlefsen, L., Bohannan, B.J.M. & Relman, D.A. The application of ecological theory toward an understanding of the human microbiome. Science 336, 1255 (2012).

16. Maier, L. & Typas, A. Systematically investigating the impact of medication on the gut microbiome. Curr Opin Microbiol 39, 128–135 (2017).

17. Van den Abbeele, P., et al. Butyrate-producing *Clostridium* cluster XIVa species specifically colonize mucins in an *in vitro* gut model. ISME J 7, 949 (2013).

18. Shah, P., et al. A microfluidics-based *in vitro* model of the gastrointestinal human–microbe interface. Nat Commun 7, 11535 (2016).

19. Kim, H.J., Li, H., Collins, J.J. & Ingber, D.E. Contributions of microbiome and mechanical deformation to intestinal bacterial overgrowth and inflammation in a human gut-on-a-chip. PNAS 113, E7 (2016).

20. Verberkmoes, N.C., et al. Shotgun metaproteomics of the human distal gut microbiota. The Isme Journal 3, 179 (2008).

21. Zhang, X., et al. Deep metaproteomics approach for the study of human microbiomes. Anal Chem 89, 9407–9415 (2017).

22. Zhang, X., et al. Metaproteomics reveals associations between microbiome and intestinal extracellular vesicle proteins in pediatric inflammatory bowel disease. Nature Communications 9, 2873 (2018).

23. Kleiner, M., et al. Assessing species biomass contributions in microbial communities via metaproteomics. Nature Communications 8, 1558 (2017).

24. Tanca, A., et al. Potential and active functions in the gut microbiota of a healthy human cohort. Microbiome 5, 79 (2017).

25. Moya, A. & Ferrer, M. Functional redundancy-induced stability of gut microbiota subjected to disturbance. Trends Microbiol. 24, 402–413 (2016).

26. Li, L., et al. Evaluating *in vitro* culture medium of gut microbiome with orthogonal experimental design and a metaproteomics approach. J Proteome Res 17, 154–163 (2018).

27. Bone, E., Tamm, A. & Hill, M. The production of urinary phenols by gut bacteria and their possible role in the causation of large bowel cancer. AJCN 29, 1448 (1976).

28. Gibson, G.R. & Wang, X. Bifidogenic properties of different types of fructo-oligosaccharides. Food Microbiol 11, 491 (1994).

29. Lesmes, U., Beards, E.J., Gibson, G.R., Tuohy, K.M. & Shimoni, E. Effects of resistant starch type III polymorphs on human colon microbiota and short chain fatty acids in human gut models. J Agric Food Chem 56, 5415 (2008).

30. Rycroft, C.E., Jones, M.R., Gibson, G.R. & Rastall, R.A. A comparative in vitro evaluation of the fermentation properties of prebiotic oligosaccharides. J Appl Microbiol 91, 878 (2001).

31. Olano-Martin, E., Mountzouris, K.C., Gibson, G.R. & Rastall, R.A. In vitro fermentability of dextran, oligodextran and maltodextrin by human gut bacteria. Br J Nutr 83, 247 (2000).

32. Saulnier, D.M.A., Gibson, G.R. & Kolida, S. *In vitro* effects of selected synbiotics on the human faecal microbiota composition. FEMS Microbiol Ecol 66, 516 (2008).

33. Zhang, X., et al. *In vitro* metabolic labeling of intestinal microbiota for quantitative metaproteomics. Anal Chem 88, 6120 (2016).

34. Turnbaugh, P.J., et al. An obesity-associated gut microbiome with increased capacity for energy harvest. Nature 444, 1027 (2006).

35. Kleiner, M., et al. Assessing species biomass contributions in microbial communities via metaproteomics. Nat Commun 8, 1558 (2017).

36. Cheng, K., et al. MetaLab: an automated pipeline for metaproteomic data analysis. Microbiome 5, 157 (2017).

37. The Human Microbiome Project, C. Structure, function and diversity of the healthy human microbiome. Nature 486, 207 (2012).

38. Zhang, X., et al. MetaPro-IQ: a universal metaproteomic approach to studying human and mouse gut microbiota. Microbiome 4, 31 (2016).

39. Liao, B., et al. iMetaLab 1.0: a web platform for metaproteomics data analysis. Bioinformatics, bty466–bty466 (2018).

40. Vidon, N., et al. Metformin in the digestive tract. Diabetes Res Clin Pract 4, 223–229 (1988).

41. Lee, H., et al. Modulation of the gut microbiota by metformin improves metabolic profiles in aged obese mice. Gut Microbes, 1–11 (2017).

42. Shin, N.R., et al. An increase in the *Akkermansia* spp. population induced by metformin treatment improves glucose homeostasis in diet-induced obese mice. Gut 63, 727 (2014).

43. de la Cuesta-Zuluaga, J., et al. Metformin is associated with higher relative abundance of mucin-degrading *Akkermansia muciniphila* and several short-chain fatty acid–producing microbiota in the gut. Diabetes Care 40, 54 (2017).

44. Plovier, H., et al. A purified membrane protein from *Akkermansia muciniphila* or the pasteurized bacterium improves metabolism in obese and diabetic mice. Nat Med 23, 107 (2016).

45. Dao, M.C., et al. *Akkermansia muciniphila* and improved metabolic health during a dietary intervention in obesity: relationship with gut microbiome richness and ecology. Gut 65, 426 (2016).

46. Zhao, L., et al. Gut bacteria selectively promoted by dietary fibers alleviate type 2 diabetes. Science 359, 1151 (2018).

47. Macfarlane, G.T., Macfarlane, S. & Gibson, G.R. Validation of a three-stage compound continuous culture system for investigating the effect of retention time on the ecology and metabolism of bacteria in the human colon. Microb. Ecol. 35, 180 (1998).

48. Charbonneau, Mark R., et al. Sialylated Milk Oligosaccharides Promote Microbiota-Dependent Growth in Models of Infant Undernutrition. Cell 164, 859–871 (2016).

49. Begley, M., Gahan, C.G.M. & Hill, C. The interaction between bacteria and bile. FEMS Microbiol Rev 29, 625–651 (2005).

50. Cao, H., et al. Secondary bile acid□induced dysbiosis promotes intestinal carcinogenesis. Int J Cancer 140, 2545–2556 (2017).

51. Ridlon, J.M., Alves, J.M., Hylemon, P.B. & Bajaj, J.S. Cirrhosis, bile acids and gut microbiota. Gut Microbes 4, 382–387 (2013).

52. Ridlon, J.M., Kang, D.-J. & Hylemon, P.B. Bile salt biotransformations by human intestinal bacteria. J Lipid Res 47, 241–259 (2006).

53. Pimentel, M., Mathur, R. & Chang, C. Gas and the Microbiome. Curr Gastroenterol Rep 15, 356 (2013).

54. Ríos-Covián, D., et al. Intestinal short chain fatty acids and their link with diet and human health. Front Microbiol 7, 185 (2016).

55. Sun, Y. & O’Riordan, M.X.D. Chapter Three – Regulation of Bacterial Pathogenesis by Intestinal Short-Chain Fatty Acids. in Advances in Applied Microbiology, Vol. 85 (eds. Sariaslani, S. & Gadd, G.M.) 93–118 (Academic Press, 2013).

56. Joly, C., et al. Impact of chronic exposure to low doses of chlorpyrifos on the intestinal microbiota in the Simulator of the Human Intestinal Microbial Ecosystem (SHIME®) and in the rat. Environ Sci Pollut Res 20, 2726–2734 (2013).

57. Wang, S. & Loreau, M. Biodiversity and ecosystem stability across scales in metacommunities. Ecol Lett 19, 510–518 (2016).

58. Amsler, C.D., Cho, M. & Matsumura, P. Multiple factors underlying the maximum motility of Escherichia coli as cultures enter post-exponential growth. J. Bacteriol. 175, 6238 (1993).

59. Relman, D.A. The human microbiome: ecosystem resilience and health. Nutr. Rev 70, S2–S9 (2012).

60. Utzschneider, K.M., Kratz, M., Damman, C.J. & Hullarg, M. Mechanisms linking the gut microbiome and glucose metabolism. J Clin Endocrinol Metab 101, 1445–1454 (2016).

61. Berney, M., Hammes, F., Bosshard, F., Weilenmann, H.-U. & Egli, T. Assessment and interpretation of bacterial viability by using the LIVE/DEAD BacLight kit in combination with flow cytometry. Appl Environ Microbiol 73, 3283–3290 (2007).

