## Supplementary figures for "An *in vitro* model maintaining taxon-specific functional activities of the gut microbiome"

**This file includes:**

**Figure S1.** Gating of live, dead and unstained bacteria according to stained gut microbiome cells, stained and heat-treated microbiome cells, and unstained microbiome.

**Figure S2.** Metaproteomic data quality.

**Figure S3.** Compositional bar chart showing species-level biomass distribution over time in the cultured microbiome of volunteers V2 and V3.

**Figure S4.** Comparison of Faecalibacterium Praunsnitzii biomass change in the MiPro- and BCM-cultured microbiomes.

**Figure S5.** Shannon-Weiner index suggesting well-maintained alpha-diversity of the microbiomes cultured from volunteers V1-3 over 48 hrs.

**Figure S6.** PCA scores plot with hierarchical clustering based on COG functional categories of microbiome proteins from volunteers V2 and V3.

**Figure S7.** Taxon-function-coupled profile in comparison with 0 hr baseline samples.

**Figure S1**

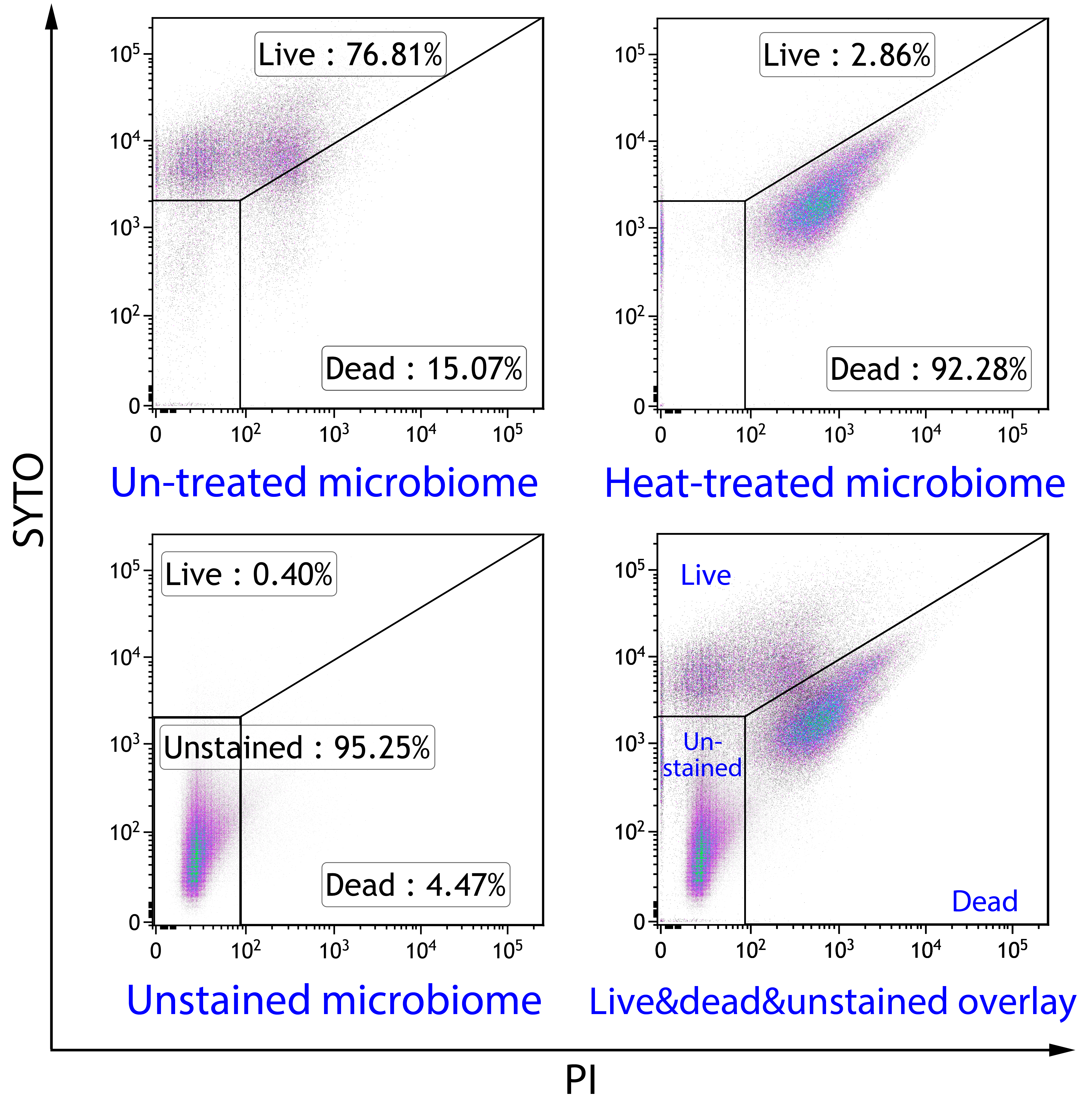

**Figure S1.** Gating of live, dead and unstained bacteria according to stained gut microbiome cells, stained and heat-treated microbiome cells, and unstained microbiome.

**Figure S2**

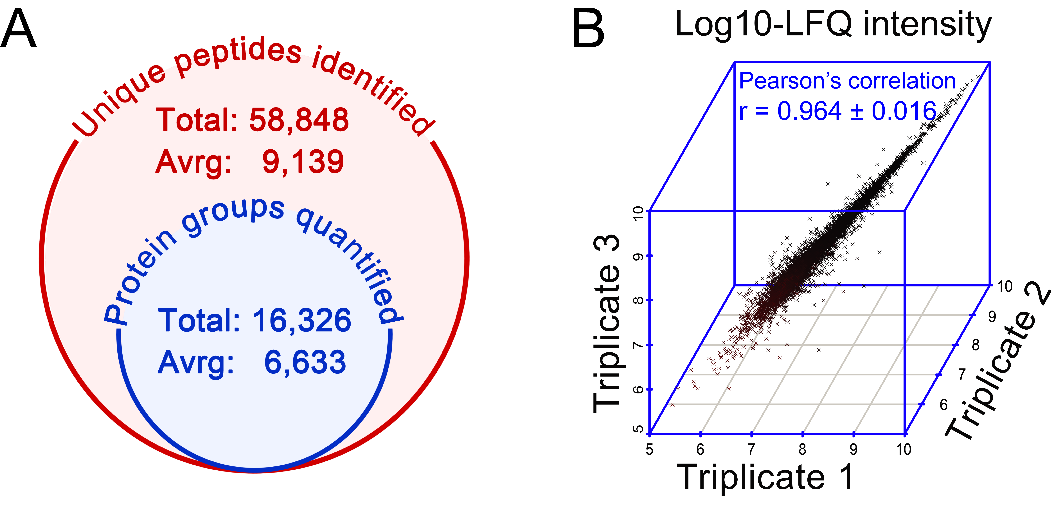

**Figure S2.** Metaproteomic data quality. (A) Venn diagram showing identification efficiency of LC-MS/MS. (B) 3D scatter plot showing metaproteomic data reproducibility of technical triplicates.

**Figure S3**

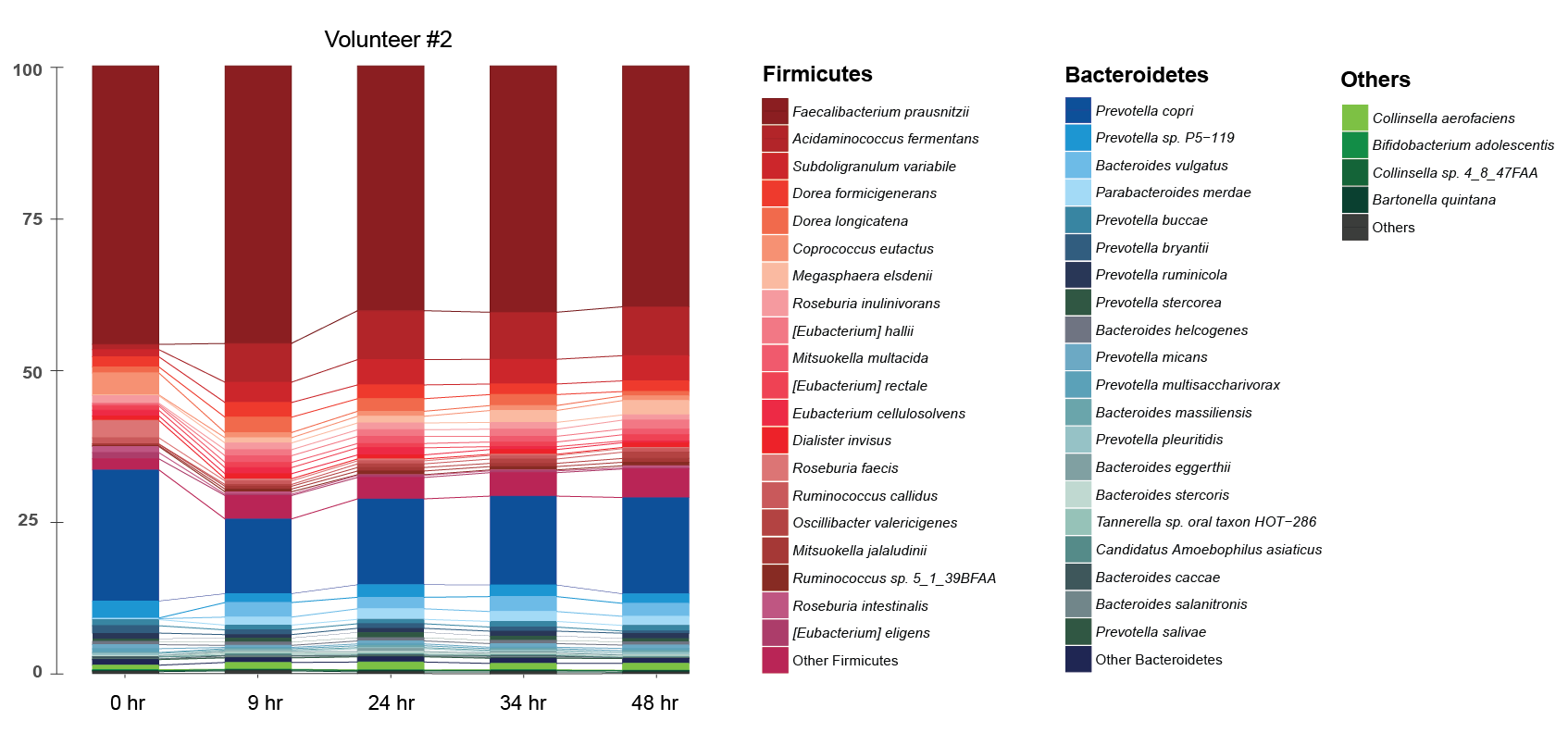

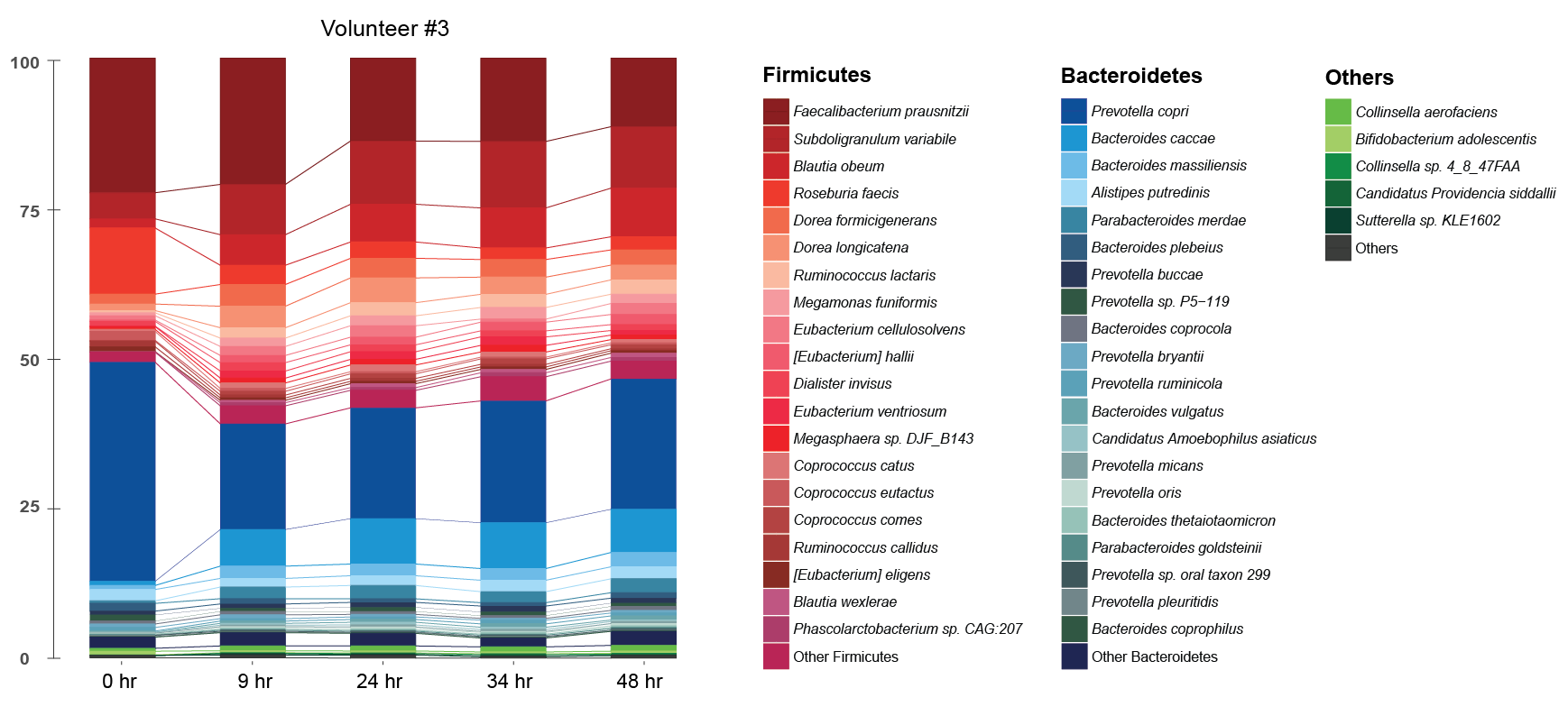

**Figure S3.** Compositional bar chart showing species-level biomass distribution over time in the cultured microbiome of volunteers V2 and V3.

**Figure S4**

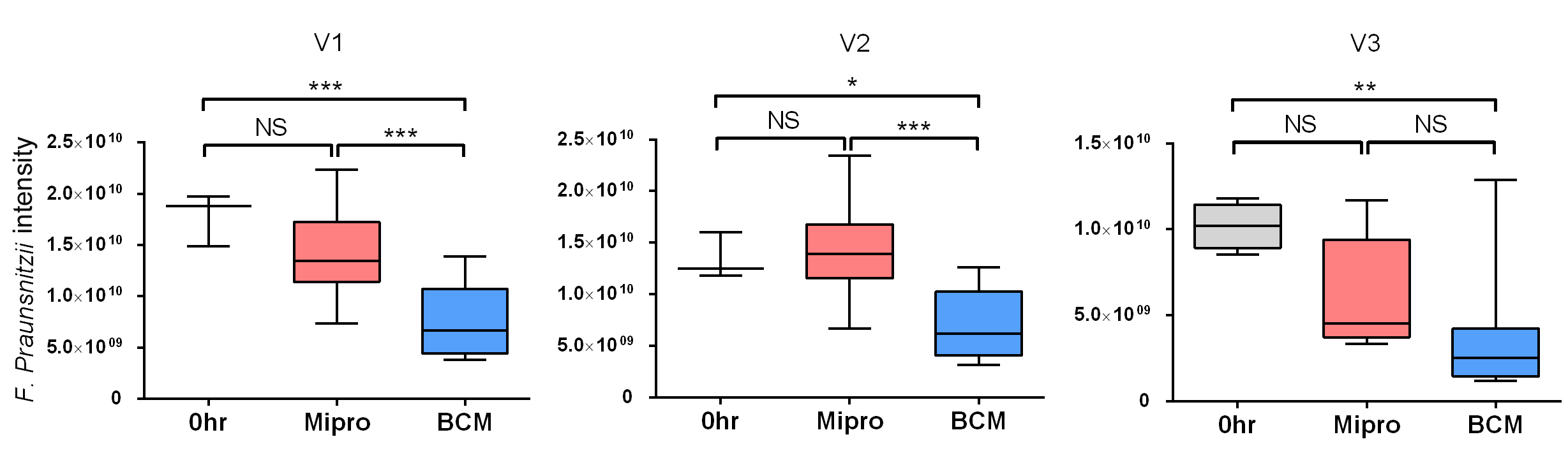

**Figure S4.** Comparison of *Faecalibacterium Praunsnitzii* biomass change in the MiPro- and BCM-cultured microbiomes. Statistical significance was evaluated by Tukey's multiple comparison test and indicated as **p*<0.05, ** *p*<0.005 and *** *p*<0.0005 .

**Figure S5**

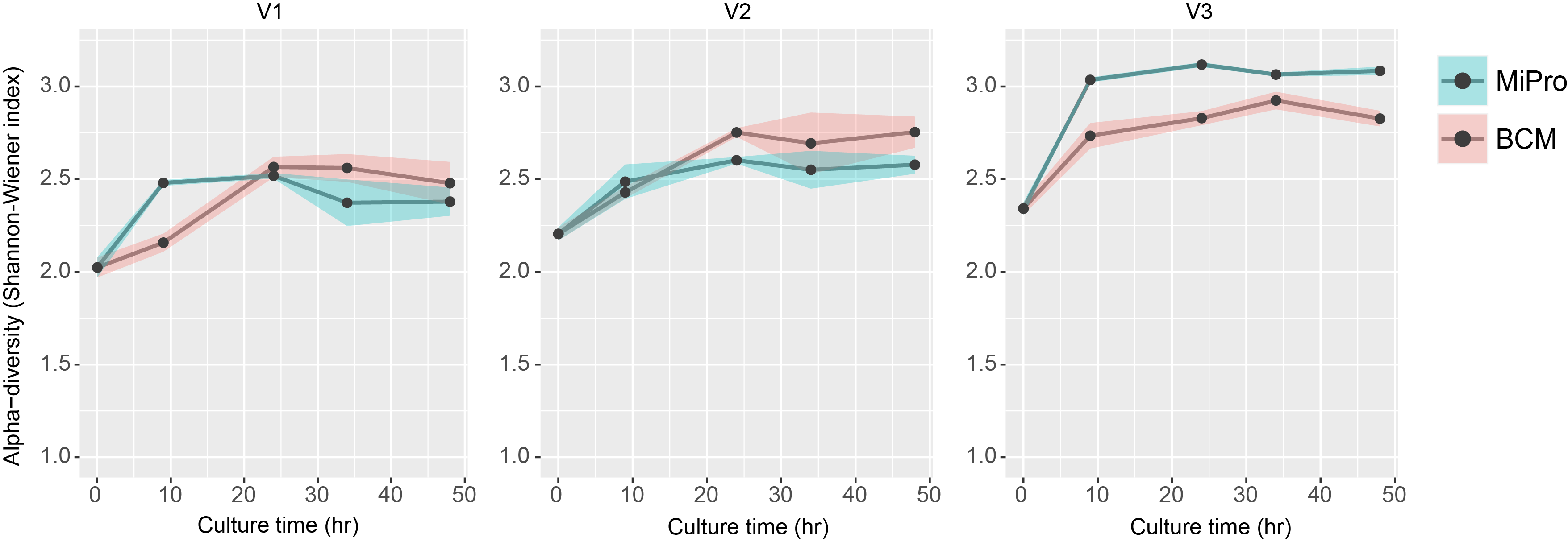

**Figure S5.** Shannon-Weiner index suggesting well-maintained alpha-diversity of the microbiomes cultured from volunteers V1-3 over 48 hr.

**Figure S6**

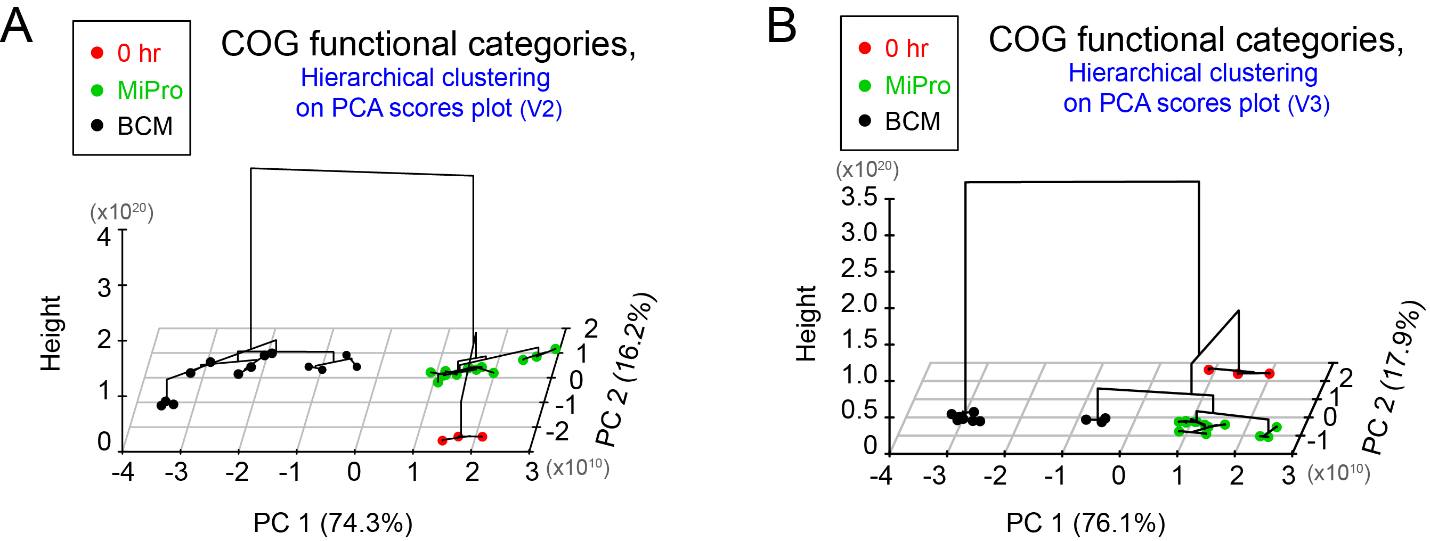

**Figure S6.** PCA scores plot with hierarchical clustering based on COG functional categories of microbiome proteins from volunteers V2 and V3. A general discrimination of 16.2%-17.9% on the PC2 axis was contributed by culture difference, whereas a larger separation on PC1 axis (74.3%-76.1%) was induced by culture in BCM medium, suggesting better functional maintenance of microbiome cultured in MiPro medium.

**Figure S7**

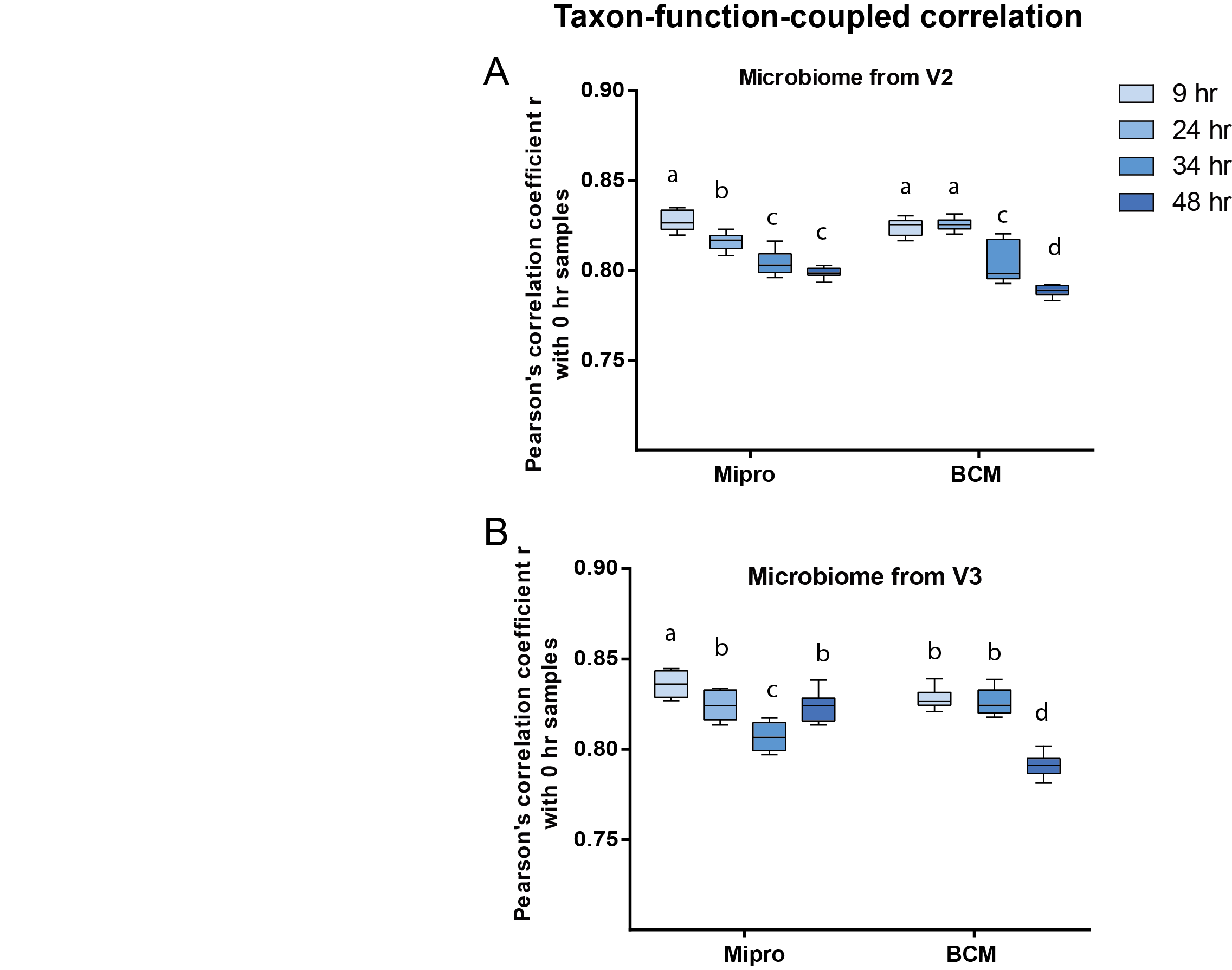

**Figure S7.** Taxon-function-coupled profile in comparison with 0 hr baseline samples. (A-B) Pearson’s correlation coefficient r of taxon-specific functional profiles between cultured and the inocula microbiomes of volunteers V2 (A) and V3 (B). Different letters indicate significant differences (*p*< 0.05) as determined by Tukey-b test.

1. ^1^ Department of Biochemistry, Microbiology and Immunology, Ottawa Institute of Systems Biology, Faculty of Medicine, University of Ottawa, Ottawa, Canada

   ^2^ Department of Statistical Sciences, Faculty of Arts and Science, University of Toronto, Toronto, Canada

   ^3^ Department of Chemistry and Biomolecular Sciences, University of Ottawa, Ottawa, Canada

   ^4^ Canadian Institute for Advanced Research, Toronto, Canada

   ^‡^ Both authors contributed equally in this work

   ^*^ correspondance,
